# Simple, static, spiking nonlinearities can account for seemingly complex interactions between visual, motor and cognitive variables in the visual cortex

**DOI:** 10.64898/2026.11.01.763479

**Authors:** Wojciech Zajkowski, Katrina Quinn, Lenka Seillier, Marie E. Bellet, Daniel A. Butts, Hendrikje Nienborg

## Abstract

Neural activity in the visual cortex is driven by the stimulus and modulated by multiple task-related variables, including spatial attention, eye movements, and perceptual decisions. These multiple influences are each typically described as nonlinear interactions such as gain or normalization, raising the question of how these diverse signals can be combined through cellular or circuit mechanisms. Here we address this problem by examining neuronal responses in macaque visual areas V2 and V3 during a visual discrimination task that dissociates spatial attention from motor planning and microsaccadic eye movements.

Using a statistical modeling framework, we show that attentional, microsaccade-related, and choice-related influences on neural activity are best explained by a simple additive stimulus-independent input modulation, which manifests as a gain via a spiking nonlinearity. In addition to explaining how interactions between variables might be accomplished mechanistically, this framework also resolves a controversy about the relationship between microsaccades and spatial attention. We find that neither microsaccade direction nor microsaccade-related neural activity accounts for attentional modulation, although these variables are often correlated, leading to apparent interactions. Thus, visual cortex can integrate sensory, motor-related, and cognitive signals through a parsimonious mechanism that preserves stable stimulus representations while supporting flexible, context-dependent modulation.

## Introduction

Neurons in the visual cortex are influenced by a wide variety of behavioral and cognitive contexts. Their responses are shaped not only by visual stimuli directly, but also by a variety of task-related factors, including spatial attention (1, 2), eye movements (3), expectations about incoming sensory input (4), and perceptual decisions (5) (Fig. 1A). As a result, their activity often reflects the combined influence of multiple variables of different origin, timescale, and functional role. Understanding how these diverse signals are integrated within sensory circuits without compromising stable stimulus representations remains a central challenge in systems neuroscience.

**Figure 1.**
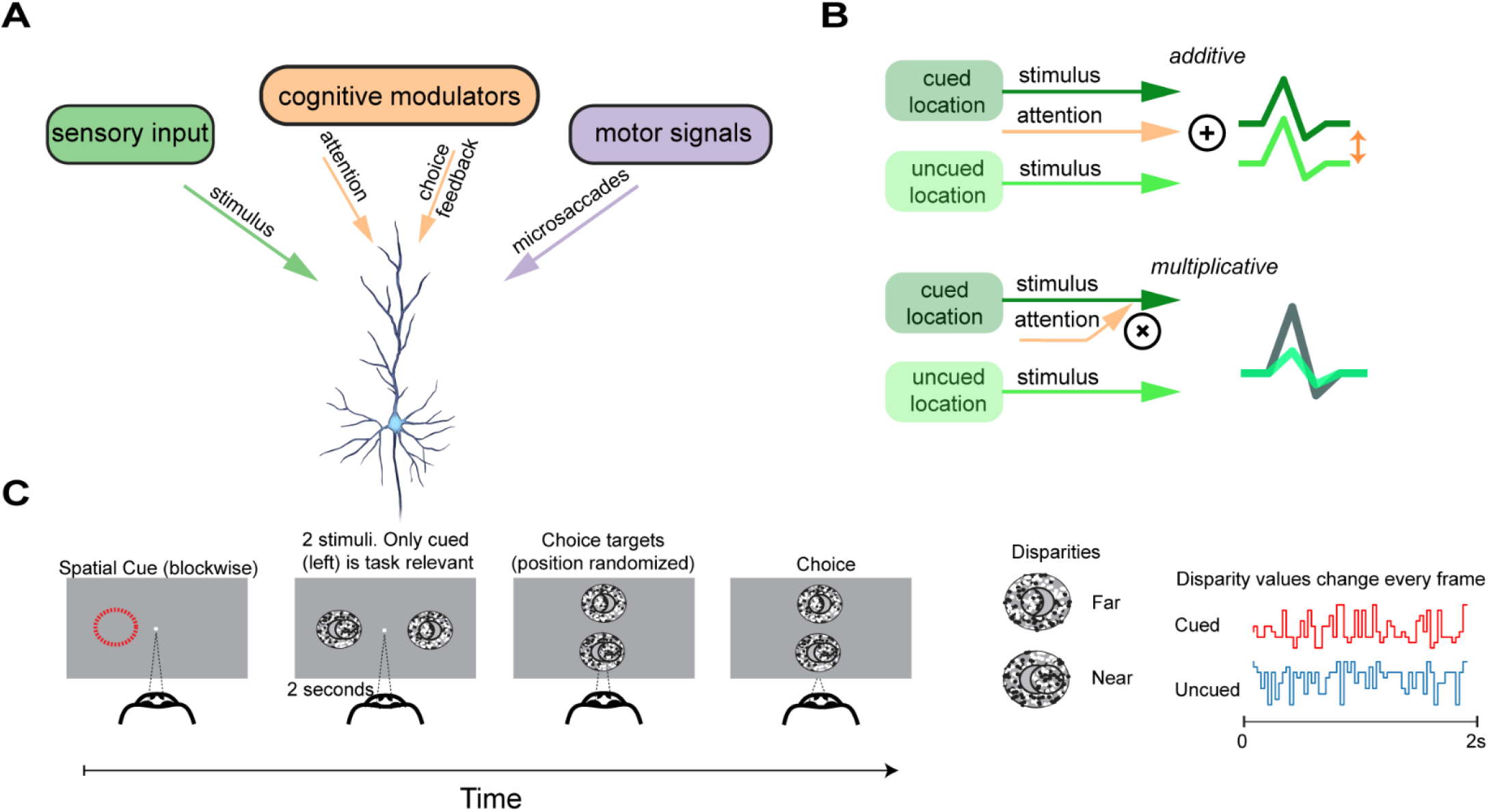
Conceptual framework and experimental task. **A.** General problem addressed in this study: responses of sensory neurons reflect not only stimulus drive but also multiple task-related influences, including cognitive modulators (e.g., attention and choice) and motor-related signals (e.g., *microsaccades*). How these signals are integrated at the level of single neurons remains unclear. **B.** Two competing accounts of cognitive modulation. In an additive model, attention and other task-related signals shift activity without altering stimulus tuning, whereas in an interactive (dependent) model, cognitive signals multiplicatively modulate stimulus-driven responses. **C.** Disparity discrimination task used to dissociate sensory input, spatial attention, and motor planning. On each trial, two stimuli were presented simultaneously, but only the cued location was task-relevant. Disparity values varied continuously over time, and behavioral choices were reported via saccades to vertically randomized choice targets, orthogonal to stimulus location.

Elucidating these effects is complicated by the fact that many task-related variables are correlated within experimental paradigms. For example, spatial attention, motor preparation, and microsaccadic eye movements often co-occur, making it difficult to disentangle their respective contributions to neural activity. Moreover, nonlinear transformations in neural responses can cause correlated inputs to appear as complex interactions at the level of firing rates, even when the underlying influences are simpler.

Here we test the hypothesis that diverse task-related influences are integrated through simple additive inputs to sensory neurons (*additive modulation*; Fig. 1B). On the surface this seems to contradict observations that the influence of cognitive factors such as attention or decisions changes the neuronal gain of stimulus-driven responses (6, 7). However, these observations are typically based on measurements of output firing rates, and they do not uniquely constrain the underlying computational mechanisms (8). In particular, gain-like effects in spiking output could arise either from multiplicative interactions at the input level, e.g. a “feature-similarity gain model” (9), tuned normalization (10–12), or alternatively, as we hypothesize here, from simpler additive inputs that are transformed by a fixed neuronal spiking nonlinearity (e.g.,(13)).

We assert that an important application of this hypothesis is to the ongoing debate over the relationship between spatial attention and microsaccadic eye movements (14–19). Some studies have suggested that attentional modulation in visual cortex depends on microsaccades directed toward attended locations, and that eye-movement-related signals gate or drive attentional effects (20–22). However, in many experimental paradigms, spatial attention, motor planning, and microsaccade direction are tightly coupled, making it difficult to dissociate their respective contributions to neural activity. This entanglement can obscure a more general question: whether cognitive variables interact directly with specific motor or sensory events, or whether they are expressed as independent, additive signals that shift the input-driven excitability. Under our hypothesis, sensory drive, motor-related signals, and cognitive feedback each affect the input-driven excitability independently, while the neuron’s fixed spike-generation nonlinearity transforms these additive inputs into the observed firing rates. As a result, complex interaction-like effects in output activity can arise without changes in stimulus tuning or specialized gating mechanisms.

To investigate this issue, we use a disparity discrimination task that decouples spatial attention, microsaccade direction, and motor planning. Our block-cueing design and orthogonal response mapping (*i.e.,* response targets being mapped to different spatial locations than category cues; Fig. 1C) allow us to dissociate attentional modulation from eye movement–related signals and stimulus-driven activity. We analyze neurons in visual areas V2 and V3, which have robust stimulus selectivity, attentional modulation and microsaccadic activity. We use a model-based approach to dissect the contributions of the individual variables despite the correlations between them.

We find the data to be best explained by additive modulation rather than changes in neural tuning curves. This demonstrates that spatial attentional modulation in V2 and V3 can be captured without invoking interactions with microsaccade-related activity or stimulus tuning at the input level. Instead, interactions in firing rates emerge from the transformation of these inputs by the spiking nonlinearity. We further find that a similar additive pattern characterizes choice-related modulation, suggesting that both attention and decision signals share a common mechanism of top-down modulation in mid-level visual cortex. This provides a parsimonious explanation for how sensory neurons can flexibly reflect behavioral context while preserving stable representations of stimulus features.

## Results

To examine how visual, motor-related, and cognitive signals jointly influence neuronal activity in mid-level visual cortex, we analyzed responses from 1,178 units in areas V2 and V3 of two macaque monkeys performing a combined spatial attention and disparity discrimination task (Fig. 1C) (23). On each trial, two random-dot stereograms were presented simultaneously, one in each hemifield, but only one stimulus was behaviorally relevant. The monkey’s task was to determine whether the central disk was protruding (‘near’) or receding (‘far’) relative to the surrounding annulus. The motor responses (*up* vs. *down*) were orthogonal to stimulus location (*left* vs. *right*), thereby decoupling spatial attention from motor planning. Moreover, the vertical locations of the choice targets were randomized from trial to trial, to uncouple the choice from the saccade direction. Behavioral results confirmed that the animals learned to selectively attend to the task-relevant stimulus while ignoring the distractor, supporting the effectiveness of the attentional manipulation (23) (Fig. S1A).

Under these task conditions, the modulation of the responses by attention was similar across the directions of microsaccades, unlike previous findings (17). Although microsaccades showed a weak bias toward attended locations (Fig. S1B-C), attentional enhancements in firing rate were comparable following microsaccades directed toward and away from the attended stimulus. Both the magnitude and time courses of the responses were similar (Fig. S1D-E), in contrast to previous reports (20, 21). This suggests that, when attention and motor planning are dissociated, microsaccades cannot account for attentional modulation, which motivated us to quantify their respective contributions via formal model comparisons.

### A statistical modeling framework dissociates independent contributions to neural activity

To quantify how stimulus drive, microsaccades, attention, and perceptual choice jointly shape neuronal responses we used a statistical modeling approach based on generalized linear models (GLMs). Each neuron’s response was generated from the sum of several components (Fig. 2A): a stimulus-driven term capturing disparity selectivity over time, a transient term capturing microsaccade-related modulation (Fig. 2B-C). Additional terms account for within-trial transients at stimulus onset and slow drifts in baseline firing across the recording session. This sum was then passed through a fixed spiking nonlinearity to generate a time-resolved predicted firing rate. This framework allowed us to test whether attention and choice were better explained by changes in stimulus tuning or microsaccade-related responses, or instead by additive modulation.

**Figure 2.**
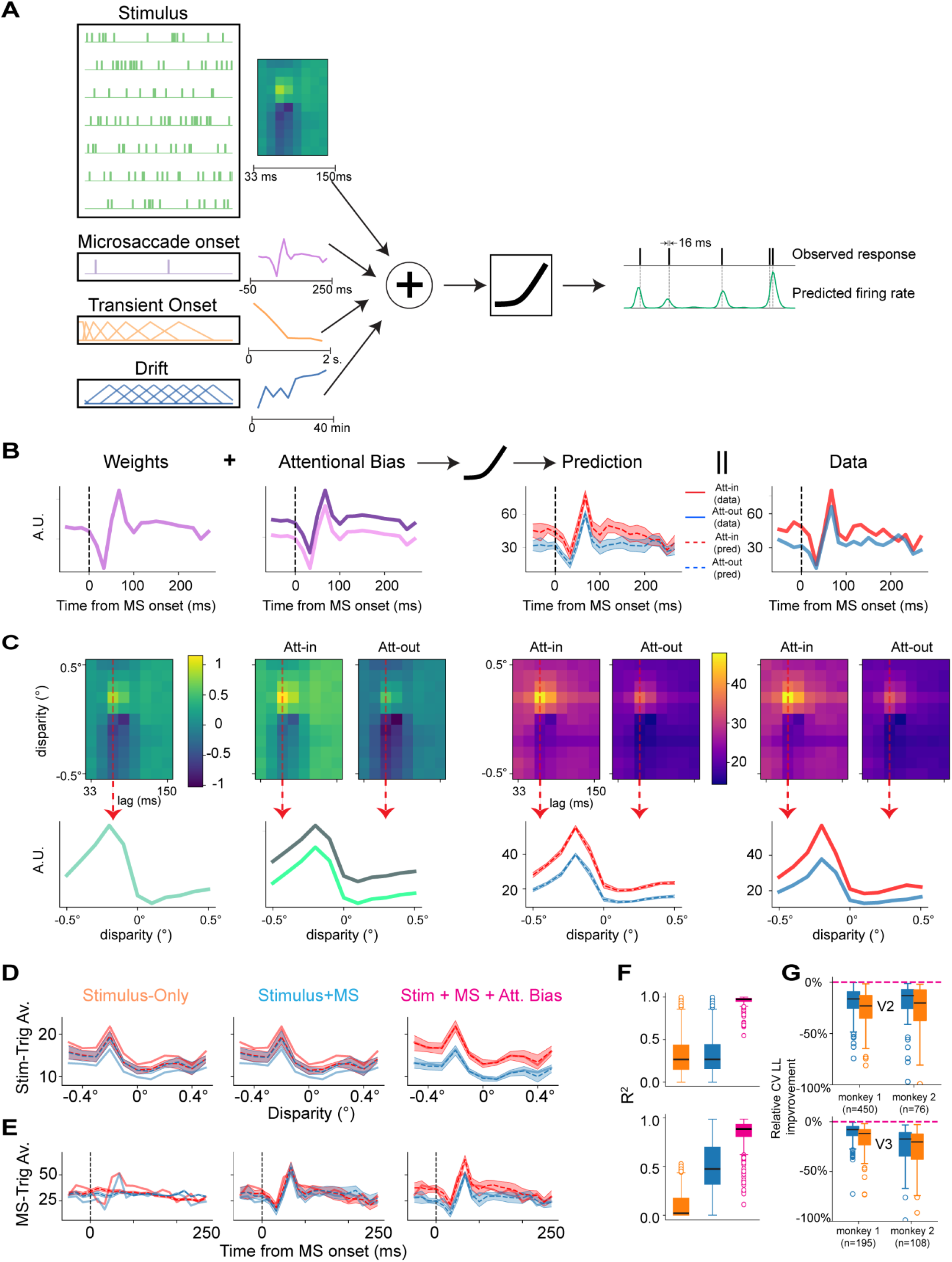
Modeling framework for dissociating sensory, motor, and cognitive influences. **A.** Schematic of model design. Separate binary regressors for each disparity and time lag describe stimulus-driven responses with high temporal resolution. Microsaccade-related activity, attentional state, slow drift, and within-trial onset transients are modeled as additional additive components. Drift and onset transients are modeled using basis functions. The second column illustrates example fitted linear weights. Right side: model output (conceptual illustration). Predicted firing rate (bottom) and observed response (top). **B.** Deriving microsaccade-related predictions from the additive modulation model (example unit). Left: fitted microsaccade kernel. Middle: additive attentional bias term. Right: predicted and observed microsaccade-triggered firing rates. **C.** Deriving stimulus tuning predictions from the additive modulation model (example unit). As in (B), but for stimulus-driven responses using a two-dimensional disparity × time-lag kernel. Bottom row shows fitted weights, model predictions, and observed responses for the most disparity-sensitive time lag. Dashed lines show mean model predictions; shaded bands indicate approximate pointwise 95% confidence intervals across cross-validation folds for both B and C. **D.** Stimulus-triggered average for both attentional conditions across 3 compared models (example unit). Only the *Stimulus + Microsaccade + Attentional Bias* (rightmost panel) can account for stimulus and attentional effects (data: solid lines; predictions: dashed lines). **E.** Microsaccade-triggered average for both attentional conditions across 3 compared models (example unit). Only the *Stimulus + Microsaccade + Attentional Bias (*rightmost panel) can account for microsaccade and attentional effects (data: solid lines; predictions: dashed lines). **F.** Variance explained for stimulus-triggered (top) and microsaccade-triggered (bottom) averages across all units. Only the additive modulation model (black box) explains both response types well. **G.** Cross-validated model comparison across all units, compared to the additive modulation model (dashed line at 0). Additive modulation model provides a significant improvement in 5-fold cross-validated fit over *Stimulus-Only* and *Stimulus+Microsaccade* models, indicating that all three factors (Stimulus, Microsaccades and Attention) are necessary to describe the features of neural activity.

We consider a model in which neural activity is described by the additive combination of stimulus tuning, microsaccade-related transients, and additive attentional modulation inside a spiking nonlinearity. We refer to this model as *additive modulation model* and compare it to more extensive models that allow for complex interactions between the stimulus, motor and cognitive variables. We confirmed that each component of the additive modulation model was necessary to account for the qualitative and quantitative features of the data: removing any one of them degraded the model’s ability to capture stimulus-driven responses, microsaccade-triggered activity, or attentional modulation (Fig. 2DE). Across the population, the additive modulation model provided a good account of neural responses and achieved the best cross-validated performance among the tested independent models (Fig 2FG; Fig S2). This additive modulation model serves as a baseline for all further comparisons. We then compare it with more flexible models that allow stimulus tuning or microsaccade-related responses to vary arbitrarily with attentional state or choice. These comparisons test whether task-related effects require such interactions or are equally well or better captured by additive, stimulus-independent shifts.

### Microsaccade-related activity is independent of attention and direction

We first asked whether microsaccade-related modulation depended on attentional state or microsaccade direction (Fig. 3A). Microsaccades produced clear modulations of neural activity in both V2 (Fig. 3B) and V3 (Fig. 3C), and the magnitude and shape of these responses could in principle vary depending on where attention was directed and whether microsaccades were directed toward or away from the attended stimulus. Such differences could reflect an interaction of microsaccade-related activity with attention or direction, rather than independent contributions of these signals.

**Figure 3.**
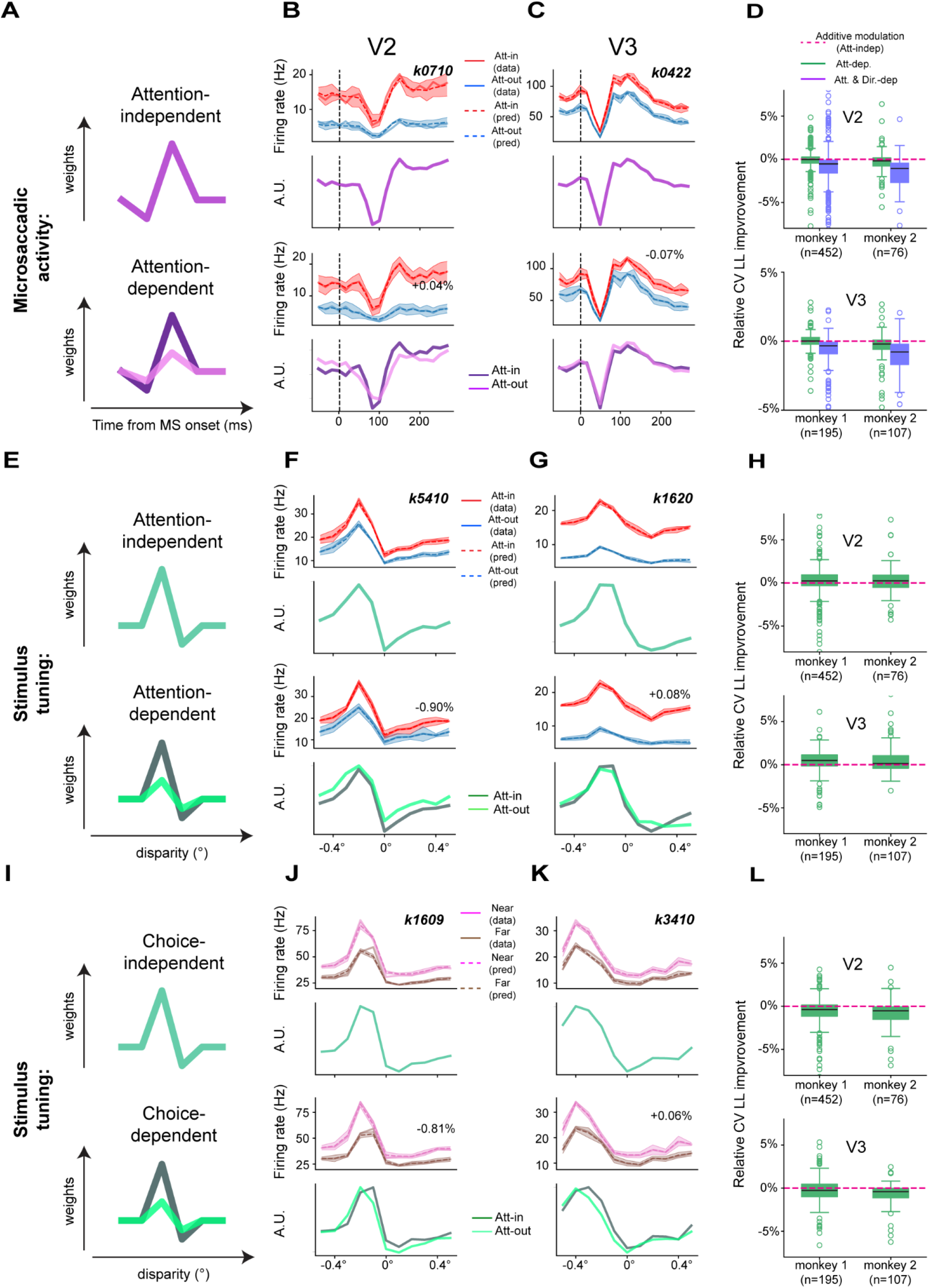
Testing whether sensory, motor-related, and cognitive signals interact or contribute independently to neural activity in mid visual cortex. **A.** Hypothesis space. Top: *Attention-independent* model, in which microsaccade-related modulation is represented by a single kernel in both attentional conditions (additive modulation model). Bottom: *Attention-dependent model*, allowing microsaccade kernels to depend on attention state. **B–C.** Example units showing that microsaccade-related activity is best described by the *Attention-independent* additive modulation model. Top and bottom panels show fits from the *Attention-independent* and *Attention-dependent* models, respectively. Solid lines show microsaccade-triggered averages; dashed lines show model fits. Percentages indicate relative improvement in cross-validated log-likelihood over the *Attention-independent* (additive modulation) model. Bottom subpanels show fitted microsaccade kernels. Dashed lines show mean model predictions; shaded bands indicate approximate pointwise 95% confidence intervals across cross-validation folds for all prediction plots. **D.** Population-level model comparison of cross-validated log-likelihoods across all units in V2 (top) and V3 (bottom). Boxes show interquartile range (IǪR); horizontal lines indicate medians. Values are expressed relative to the additive modulation model (dashed line at 0). **E.** Hypothesis space. Top: *Attention-independent* model with a single stimulus kernel shared across conditions (additive modulation model). Bottom: Attention-dependent model allowing separate stimulus kernels for attended and unattended conditions. **F-G.** Example units showing that stimulus disparity tuning is best described by the Attention-independent model. Top and bottom panels show fits from the Attention-independent and Attention-dependent models, respectively. Top subpanels display stimulus-triggered averages (solid lines) and model fits (dashed lines) across disparity values. Percentages indicate improvement over the *Attention-independent* additive modulation model. Bottom panels show fitted disparity kernels. **H.** Cross-validated model comparison across all units. Allowing attention-dependent stimulus tuning provides no systematic improvement over the *Attention-independent* additive modulation model (black dashed line). Box plots show medians and IǪRs. **I.** Hypothesis space. Top: Choice-independent model with a single stimulus kernel shared across choices (additive modulation model). Bottom: Choice-dependent model allowing separate stimulus kernels for each choice category. **J–K.** Example units showing that disparity tuning is best described by the *Choice-independent* model. Top and bottom panels show fits from the *Choice-independent* and *Choice-dependent* models, respectively. Top subpanels show observed (solid) and predicted (dashed) firing rates. Bottom panels show fitted stimulus kernels across disparity values. Percentages indicate improvement over the *Choice-independent* additive modulation model. **L.** Cross-validated log-likelihood comparison across all units. The *Choice-dependent* model does not improve predictive performance relative to the *Choice-independent* additive modulation model (dashed line). Box plots show medians and IǪRs.

To distinguish these possibilities, we compared the additive modulation model, in which a single microsaccade kernel is shared across conditions and attention contributes independently, with more flexible models that allowed microsaccade-related activity to depend on attention and direction. In the *Attention-dependent* model, separate microsaccade kernels were fit for attended and unattended conditions (Fig. 3A). In the *Attention × Direction-dependent* model, these kernels could additionally vary with microsaccade direction (toward vs. away), yielding four condition-specific microsaccade kernels.

At the level of single units, the additive modulation model provided good predictions of microsaccade-triggered activity across attention conditions. Microsaccade kernels estimated separately for attended and unattended trials in the *Attention-dependent* model were very similar, with median kernel similarities (R^2^; see: *Methods*) of 0.70 (SE = 0.01) in V2 and 0.72 (SE = 0.02) in V3 (Fig.3B,C). Likewise, allowing microsaccade-related responses to depend jointly on attention and direction (*Attention × Direction-dependent* model) produced little qualitative change in the fitted kernels: median similarities between kernels for microsaccades directed *toward* versus *away* from the attended stimulus ranged from 0.59 (SE = 0.02) to 0.62 (SE = 0.01) across areas and attention conditions.

Population-level model comparisons confirmed these observations, indicating that neural activity in V2 and V3 is well described by a single, attention-invariant kernel, with an additive attention-based baseline shift. Across all tested units, neither the *Attention-Dependen*t nor the *Attention × Direction-Dependent* model outperformed the additive modulation model based on 5-fold cross-validated log-likelihood scores (Fig. 3D). In fact, both models slightly decreased predictive performance, indicating that the additional parameters did not improve the model predictions but rather led to overfitting. Specifically, the *Attention-dependent* model showed an average performance decrease of 0.5% relative to the additive modulation model (ranging from 14.2% decrease to 6.4% increase, with median = 0.1% decrease). The *Attention × Direction-dependent* model performed even worse, with an average decrease of 1.7% (ranging from 50.3% decline to 10.9% increase, median = 1.0% decrease).

### Attentional modulation preserves stimulus tuning

Next, we asked whether attentional modulation interacted with stimulus disparity. At the level of firing rates, attention produced disparity-dependent changes in response magnitude, consistent with the gain-like modulation of tuning curves reported previously (Fig. 3FG). We therefore asked whether this apparent interaction reflected an attention-dependent change in the underlying stimulus tuning, or whether it could instead emerge from additive attentional modulation. To distinguish these possibilities, we compared the additive modulation model, which assumes a single stimulus kernel shared across attention conditions, with a more flexible *Attention-dependent* tuning model in which separate stimulus kernels were fit for attended and unattended trials (Fig. 3E).

At the level of single units, disparity tuning was well described by the shared-kernel additive modulation model. Allowing the stimulus tuning to vary with attention produced little qualitative change in predicted responses, and disparity kernels estimated separately for attended and unattended conditions showed substantial overlap in both shape and temporal profile: across units, the median similarity (R^2^) between them reached 0.86 (SE = 0.02) in V2 and 0.88 (SE = 0.09) in V3 (Fig. 3F-G).

Population-level model comparisons confirmed that attentional modulation in V2 and V3 was largely stimulus-independent. The *Attention-dependent* stimulus tuning model provided no systematic improvement in cross-validated performance relative to the additive modulation model, with a median increase of +0.3% and substantial variability across units. In many cases, the additional parameters led to slight decreases in predictive accuracy, consistent with overfitting (Fig. 3H). Together, these results show that the gain modulation of the tuning curve by attention was well described by a simple additive shift in the model at the input level. How do independent, additive inputs lead to these interactions between stimulus tuning and attention?

### Additive input modulation gives rise to multiplicative effects through spiking nonlinearity

In our additive modulation model, the summed linear contributions of stimulus drive, microsaccades and attention, are transformed into firing rates by a static spiking nonlinearity. Because this nonlinearity is rectifying and expansive near the spiking threshold, additive modulation in the linear input domain increases the slope of the input–output relationship, producing proportional scaling of responses across stimulus conditions. As a result, uniform baseline shifts can manifest as gain-like changes in firing rate without requiring multiplicative interactions or stimulus-specific modulation (Fig. 4A–D).

**Figure 4.**
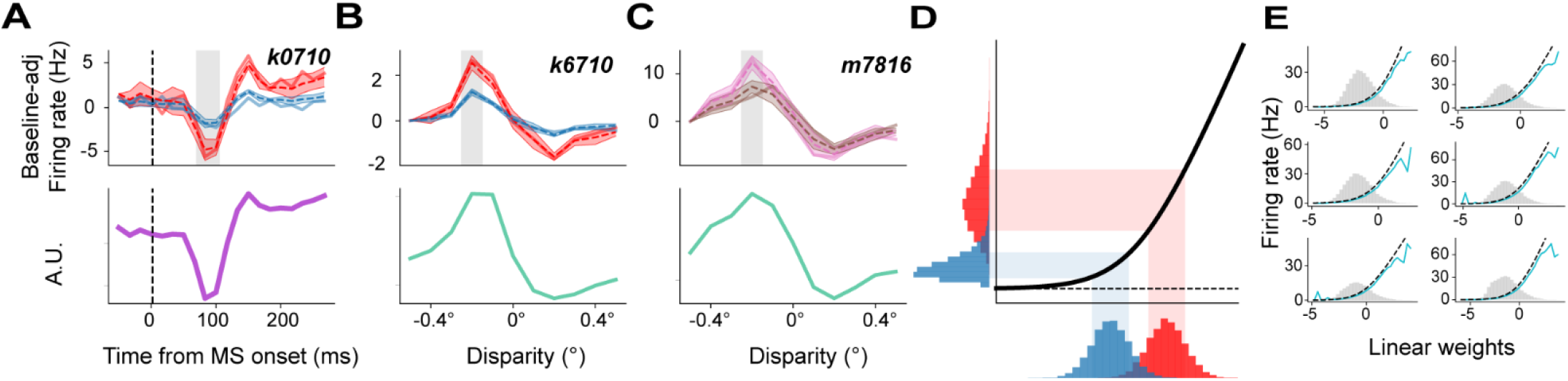
Spiking nonlinearity transforms additive inputs into gain-like effects. **A–C.** Example units illustrating apparent multiplicative modulation arising from the additive modulation model. Panels show examples of attentional modulation of microsaccade-related activity (A), attentional modulation of stimulus tuning (B), and choice-related modulation of stimulus tuning (C). Shaded regions highlight response ranges for which output firing rates appear multiplicatively scaled despite additive inputs. Dashed lines show mean model predictions; shaded bands indicate approximate pointwise 95% confidence intervals across cross-validation folds. **D.** Conceptual illustration showing how the *so*ftplus nonlinearity (black curve) transforms additive shifts in linear input (distributions below x-axis) into multiplicative-like changes in output firing rate (distributions along y-axis). **E.** Empirical fits of the *so*ftplus nonlinearity for six example units. Gray histograms show the distribution of linear predictions; curves show the relationship between predicted input and observed firing rates.

This transformation provides a simple explanation for why attentional effects appear multiplicative when assessed at the level of firing rates. For individual units, the additive modulation model accurately reproduced response patterns that resemble classic gain modulation, including proportional increases in firing across disparity values and enhanced responses during microsaccade-triggered activity (Fig. 4A–C). Empirical fits of the spiking nonlinearity to example units further confirm that this transformation captured the observed relationship between linear predictions and firing rates across a wide dynamic range (Fig. 4E).

### Choice-related modulation also preserves stimulus tuning

To test whether this explanation held more generally, we next examined whether it also applied to the modulation of spiking activity associated with perceptual choice. Previous work has shown that visual sensory neurons can exhibit choice-related activity (5, 24), at least partially explained by feedback (7, 23) raising the possibility that decision-related signals interact with sensory representations. Indeed, previous work identified systematic gain changes of the stimulus tuning curve with choice (7). If our explanation above holds more generally, such choice-related gain change of the stimulus tuning might also be described by a simple additive shift related to choice. To test this possibility, we compared the additive modulation model with an added choice-bias and attention parameters (c*hoice-independent* model) to a more flexible *Choice-dependent* tuning model in which separate stimulus kernels were fit for trials associated with “*near*” and “*far*” choices (Fig. 3I).

At the level of single units, disparity tuning was indeed well described by a single, choice-independent stimulus kernel for both the attended and unattended stimulus conditions. Allowing stimulus tuning to depend on choice and attention produced little qualitative change in predicted responses, and disparity kernels estimated separately for near and far choices showed substantial overlap (Fig. 3J-K). Across units, the median similarity (R^2^) between near- and far-choice disparity kernels was 0.82 (SE = 0.07) in V2 and 0.86 (SE = 0.08) in V3, indicating highly similar tuning across choice conditions.

Population-level model comparisons confirmed that choice-related modulation in V2 and V3 did not require changes in disparity tuning to explain observed firing-rate differences. The *Choice-dependent* tuning model did not improve cross-validated performance relative to the additive modulation model; instead, it produced a small median decrease in predictive accuracy (−0.4%), with substantial variability across units (Fig. 3L).

## Discussion

Our results show that task-related influences on neuronal activity in macaque V2 and V3, including spatial attention and choice feedback, are well described by a single straightforward mechanism. Across analyses, factors that modulate neural responses, such as cognitive effects like attention or decision-related feedback, act primarily through additive modulation, while leaving stimulus tuning largely unchanged. Similarly, while microsaccadic activity typically had a characteristic temporal profile, it could simply be added at the input level and was independent of stimulus tuning and cognitive effects. When combined with a fixed spiking nonlinearity, these additive inputs gave rise to the multiplicative, gain-like effects that are commonly observed at the level of firing rates.

Our findings also clarify the relationship between spatial attention and microsaccades in mid-level visual cortex, showing that attentional modulation is not explained by microsaccadic eye movements, and that motor-related and cognitive signals can be integrated without direct interaction. Earlier studies in V4 proposed that attention-related neural enhancement emerges only after a microsaccade directed toward the attended stimulus, implying that microsaccades drive attentional effects (20, 21). More recent work in the superior colliculus and human EEG has challenged this causal view, showing that microsaccades instead track ongoing fluctuations in attention without directly generating them (16, 25, 26). Here, by using a task design that dissociates attention from motor planning, we show that attentional modulation in V2 and V3 is not explained by microsaccade direction or by interactions between attention and eye-movement– related signals. Instead, microsaccades contribute transient modulations that are largely invariant across attentional conditions, while attentional effects manifest independently as baseline shifts.

Together, these results suggest that microsaccades are best viewed not as drivers of attentional modulation in early visual cortex, but as one of several task-related signals whose effect on neural activity need not require direct interactions. More broadly, these results raise the question of whether attention itself operates through mechanisms distinct from other forms of cognitive influence, or whether different cognitive signals share a common mode of action in early sensory areas. Our analyses suggest the latter, showing that attention and choice modulate neuronal responses in similar ways, acting through stimulus-independent baseline shifts rather than changes in sensory tuning.

Our findings build on earlier work showing that nonlinear transformations can convert additive inputs into gain-like changes in neuronal responses, both in biophysical models and in exponential-link GLMs (13, 27). Normalization models likewise account for attentional modulation through nonlinear computations, with the resulting effects depending on the spatial and feature selectivity of the stimulus, attentional field, and normalization pool (10, 11, 28, 29). Our results extend this general principle to cognitive influences in V2 and V3, showing that spatial attention and decision-related feedback can be captured by stimulus-independent additive inputs transformed by a fixed output nonlinearity, without requiring changes in stimulus tuning.

A plausible physiological substrate of baseline shifts involves the activity of neuromodulatory systems, such as cholinergic, noradrenergic and dopaminergic projections, which can modulate cortical excitability in response to salient events on relatively slow timescales (30, 31). Despite their diffuse projections, neuromodulators can produce functionally specific attentional effects. Acetylcholine and dopamine can selectively enhance the representation of attended stimuli while suppressing distractors (32–35). This apparent paradox in which a broadly distributed neuromodulatory signal produces spatially or feature-specific effects may be resolved if these neuromodulators generate shifts in neuronal excitability that are subsequently converted into multiplicative changes in output through the nonlinear properties of spike generation (36). This mechanism would reconcile the slow, widespread action of neuromodulators with the rapid, spatially selective attentional effects observed electrophysiologically, offering a biologically plausible implementation of cognitive control. Directly testing this hypothesis by linking specific neuromodulatory inputs to baseline shifts in targeted cortical circuits remains a promising direction for future work.

Nonlinear transformations have long been recognized as critical determinants of how sensory signals are represented in neural firing. In V1, for example, spike threshold can amplify gradual differences in subthreshold responses, contributing to making neurons appear as two classes: simple and complex cells (37). Related nonlinear operations are central to energy-model accounts of visual feature extraction: squaring and pooling the outputs of quadrature filters generates phase-invariant motion signals (38), and analogous operations can produce binocular disparity selectivity (39). Nonlinear response thresholds also contribute substantially to decorrelation and efficient information transmission in retinal ganglion cells (40). Our findings extend this principle to cognitive modulation, showing that nonlinearities may determine not only what sensory information neurons encode, but also how cognitive signals are represented in midlevel visual cortex.

It is likely that the balance between additive and interaction-based modulation varies across the cortical hierarchy and task demands. In higher visual areas such as V4, or in oculomotor structures like the superior colliculus, attentional and motor-related signals may be more tightly coupled to specific stimulus features or movement plans, consistent with evidence linking attentional modulation to oculomotor processing in these regions (20, 41–43). Importantly, additive modulation at one processing stage need not remain additive at subsequent stages. Once transformed by a spiking nonlinearity, an additive input can produce stimulus-dependent, gain-like changes in neuronal output; downstream areas receiving this output may therefore inherit an already modulated sensory representation that is better described by interactions between cognitive state and stimulus drive. This provides one possible explanation for why interactions between attention, eye movements, and sensory encoding may be more prominent in higher visual or oculomotor areas, consistent with reports of presaccadic enhancement and direction-specific modulation (16, 20, 21). Our results do not rule out such mechanisms but show that they are not required to explain attentional and decision-related modulation in mid-level visual cortex.

Moreover, our analyses are focused on firing-rate modulation and are therefore agnostic to other dimensions of neural activity, including correlated variability (44, 45), oscillatory synchronization (46), or subthreshold dynamics (47), which may also be influenced by attention or perceptual choice. Together, these considerations suggest that additive baseline modulation represents a fundamental, but not exclusive, mode of top-down influence in sensory cortex. Identifying how this mechanism interacts with more specialized or context-dependent forms of modulation across cortical hierarchy will be an important goal for future work.

## Methods

### Dataset

We analyzed a previously published dataset (23) using less stringent behavioral inclusion criteria (for details see *Supplementary materials*). Only completed trials were included. Units with pronounced nonstationarity in firing rate or insufficient attentional modulation (attention index < 0.1) were excluded, resulting in the removal of 428 of 1,191 V2 units (35.9%) and 356 of 880 V3 units (40.5%). For analyses requiring attentional modulation, we further excluded units whose attention index did not exceed 0.1. This removed an additional 235 of the remaining 763 V2 units (30.8%) and 222 of the remaining 524 V3 units (42.4%), leaving 528 V2 and 302 V3 units.

### Animals and task

Two adult male rhesus macaques (*Macaca mulatta*; Monkey M1: 7 kg; Monkey M2: 9 kg; both aged 13 years; pair-housed) with surgically implanted titanium head post and bilateral titanium recording chambers positioned over the opercular region of area V1 performed a cued disparity discrimination task.

On each trial, two dynamic random-dot stereograms were presented simultaneously, one in each hemifield, and animals reported whether the cued stimulus appeared nearer or farther than its surround *via* a saccade to one of two choice targets. At the beginning of each attention block (50 trials), the relevant hemifield was indicated by three initial trials presenting a single stimulus on that side. Once the animal fixated at a central point, two dynamic random-dot stereograms were presented simultaneously for 2s, one in each hemifield. The stimuli were statistically matched but independently generated, and only the cued stimulus provided information relevant to the correct decision. The monkeys’ task was to judge whether the central disk within the cued stimulus appeared to protrude (“*near*”) or recede (“*far*”) relative to its surround. Following stimulus presentation, two choice icons (each representing a “*near*” or “*far*” choice with 100% disparity signal) appeared, typically offset ∼1–3° horizontally toward the cued side. Their vertical position (3°–4° above and below fixation) varied randomly across trials, preventing the animals from anticipating the saccade direction during stimulus viewing. Decisions were reported *via* a saccade to one of the icons, and correct choices were rewarded with a liquid reward. Details about experimental procedure and visual stimulus design can be found in *Supplementary materials*.

### Electrophysiological recordings

Extracellular single and multi-unit activity was recorded from areas V2 and V3 using multi-channel laminar probes (Plexon, TX, USA; V/S Probes, 24/32 channels, 50–100 μm inter-contact spacing). Because recording sites in V3 and V3A could not be reliably distinguished, we pooled recordings from these areas and refer to them collectively as V3 throughout. The probe’s position was adjusted between sessions to minimize the likelihood of sampling the same units across sessions. Eye movements were monitored binocularly with the Eyelink 1000 (SR Research, OTT, Canada) at a sampling rate of 500 Hz. Neuronal signals were acquired using Trellis (Ripple Neuro, UT, USA) interfaced with Matlab (R2014b, Mathworks) through Xippmex (v1.2.1), amplified, digitized, and filtered (250 Hz to 5 kHz) with the Ripple Grapevine System (Ripple Neuro, UT, USA).

### Microsaccade detection and direction

Microsaccades were detected using a neural network–based algorithm (48). Microsaccade direction was determined from the change in eye position between pre- and post-microsaccade samples and classified as left, right, up, or down based on the resulting movement angle.

### Modelling

We analyze the data using a statistical modelling approach based on Generalized Linear Model framework. This modelling framework was chosen over traditional spike-triggered average methods because it allows us to distinguish between independent effects of visual stimuli, microsaccade occurrences, and attentional states, and because it can handle nonlinear response properties (49, 50).

#### Nonlinearity

We used a *so*ftplus nonlinearity as the link function: F(z) = ln(1 + exp(z)). This choice ensures non-negative firing rates, approximates rectification, and maintains concavity in the log- likelihood surface for efficient optimization. Unlike exponential links, *so*ftplus provides a smooth transition from subthreshold to suprathreshold regimes, better capturing graded responses in V2/V3 neurons under varying attention condition and microsaccade events.

We assume that the spiking rate *r* at time *t* follows a Poisson distribution with mean *λ*(*t*), given by a *so*ftplus nonlinearity applied to a linear combination of predictors:

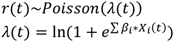

where *X_i_*(*t*) denotes the *i*-th predictor time series, and *β_i_* are the corresponding regression coefficients.

#### Stimulus

Stimulus effects were modeled with finite-impulse-response kernels (independent coefficients at each disparity *d* and time lag *τ*):

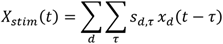

where *s_d_*,*_τ_* are the linear kernel coefficients for disparity *d* and time-lag *τ* (from 33 to 150 ms after stimulus onset), and *x_d_*(*t*) is a binary indicator that equals 1 when disparity *d* was presented at time *t* and 0 otherwise. In attention-independent stimulus models, a single stimulus kernel *s*(*d*, *τ*) was fit. In attention-dependent models, separate stimulus kernels were fit for the two attentional conditions.

#### Microsaccades

Microsaccade effects were modeled using a finite-impulse-response kernel with independent coefficients at each time lag *τ*:

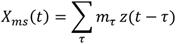

where *m_τ_* are the linear kernel coefficients at time lag *τ, (*from -50 to 250 ms relative to microsaccade onset) and *z*(*t*) is a microsaccade event indicator that equals 1 at microsaccade onset times and 0 otherwise. Microsaccade effects were modeled using one (attention-independent), two (attention-dependent), or four (attention- and direction-dependent) kernels.

#### Attention and choice

Attentional state and behavioral choice were modeled either as additive binary predictors of (attention: attended stimulus inside vs. outside the receptive field; choice: near vs. far), or by allowing separate stimulus kernels for each condition, permitting these factors to modulate stimulus tuning.

#### Transient Onset

Slow changes in firing rate within a trial were modeled using eight tent basis functions spanning the trial duration, with anchor points more densely spaced early in the trial to capture onset transients.

#### Drift

Session-level nonstationarities were modeled using tent basis functions spanning the experimental session, with anchor points placed at regular intervals (one per two attention blocks), allowing for smooth firing-rate drift over time.

#### Model Fitting

Parameters were estimated *via* maximum a posteriori (MAP) approach (50, 51). Regularization (L2 penalty on coefficients) prevented overfitting, with hyperparameters tuned via 5-fold cross-validation (50). We assessed the goodness of fit by using 5-fold cross-validation *(i.e.,* 80% of the data was used to fit model parameters, and the remaining 20% served as a testing set.

#### Model comparison

To compare models, we calculated log-likelihood (LL) scores for each model type for a given unit. We then calculated model improvement over the *drift-only* model as a difference in LL scores of the drift-only and other model types (higher values represent greater improvement). This score reflected the relative improvement in fit over the *drift-only* (null) model. When comparing non-null models we divided all relative LL scores by the LL score of the *Independent* model, to get a score reflecting relative improvement score (e.g., value of 1.05 would indicate a 5% improvement in fit over the *Independent* model).

### Kernel similarity analysis

We quantified kernel similarity using a variance-explained measure (R^2^) adjusted for variability across cross-validation folds. For each unit, we computed the mean squared difference between condition-specific kernels after mean-centering each fold, then corrected this between-condition difference by subtracting the average fold-to-fold mean squared difference observed within each condition. The resulting adjusted R^2^ therefore measures similarity between condition-specific kernel shapes after accounting for variability expected from cross-validation folds alone.

## Acknowledgments

This work is supported by the National Eye Institute Intramural Research Program at the National Institutes of Health (1ZIAEY0000570-1). The contributions of the NIH authors were made as part of their official duties as NIH federal employees, are in compliance with agency policy requirements, and are considered Works of the United States Government. However, the findings and conclusions presented in this paper are those of the author(s) and do not necessarily reflect the views of the NIH or the U.S. Department of Health and Human Services. D.A.B. is supported by the National Science Foundation (NSF IIS1350990) and the National Institutes of Health NIH R01 EY037347

## Supplementary Methods

### Inclusion criteria

We included completed, non-instruction trials on which the animal maintained fixation and registered a response, including both correct and incorrect responses (54,508 trials across 81 sessions). Units were required to have a mean firing rate exceeding 4 spikes/s on zero-signal trials (total of 2073 out of 2595 units). For each unit, periods of pronounced non-stationarity were removed. To do so we computed a running 20-trial average spike count, calculated separately for each attention condition. Periods for which this running average decreased below 20% of its peak were removed, and the longest continuous segment of included trials was used for further analysis. Additionally, we removed units based on mean and variance stationarity of the signal. For each unit, the session was split into three parts. For each of those parts, the signal mean and variance were calculated. We then calculated the proportion of the highest to lowest value for each of those measures. If this ratio exceeded 1.5 for either the mean or variance, the unit was removed from further analysis. This allowed us to remove units with highly nonstationary signal profiles. Based on these criteria, we excluded 429/1192 V2 units (36.0%) and 355/881 V3 units (40.3%).Finally, since we sought to characterize the attentional modulation, for the model comparisons, we also excluded units with attention index below 0.1. The attention index (calculated per unit) was formalized as:

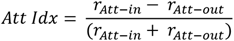

where *r* is the average firing rate on each 2 sec trial and *Att-in* and *Att-out* refer to the attentional conditions (i.e., whether the attended stimulus was within the receptive field of the unit) across all disparity values. Based on the attention criterium, additional 235 V2 units (30.8%) and 222 V3 units (42.2%) were excluded. The dataset that was further analyzed in detail consisted of (450 V2, 195 V3/V3a units from M1 and 76 V2, 108 V3/V3a units from M2).

### Procedure

We first pinpointed recording locations using individual tungsten-in-glass electrodes (Alpha Omega, Nazareth, Israel), choosing sites that showed a preference for visual disparity and had receptive fields in the desired locations. These electrodes were carefully inserted into areas V2 and/or V3/V3a. To ensure accuracy, we used anatomical MRI scans of the animals’ brains to guide the probes, which were advanced roughly perpendicular to the cortical surface with a microdrive system (NaN Instruments, Israel) and custom-designed holders. We identified V2 following established procedures (Nienborg and Cumming, 2006) and later confirmed its location by observing a change in receptive field position and size relative to V1. We located V3/V3a by noting further shifts and enlargements of receptive fields compared to V2, along with a grouping of responses based on binocular disparity (Hubel et al., 2015).

### Stimuli

Visual stimuli were presented on a screen using a DLP LED Propixx projector (VPixx, Saint-Bruno, Canada) with a resolution of 1920 × 1080 pixels. The screen had a mean luminance of 30 cd/m2 and used linearized gray values. It operated at 100 Hz or 60 Hz for each eye, in conjunction with an active circular polarizer (DepthǪ, Lightspeed Design Inc., WA, USA) running at 200 Hz or 120 Hz. Monkeys viewed the screen binocularly through passive circular polarizing filters. Viewing distances were 103.0 cm for monkey A and 97.5 cm for monkey B. Stimuli were created in Matlab (Mathworks) using custom code based on existing research (Eastman and Huk, 2012) and the Psychophysics toolbox (Kleiner et al., 2007).

The stimuli themselves were circular dynamic random-dot stereograms (RDS), composed of 50% black and 50% white dots, typically with a radius of 0.08° and a dot density of 50%. Each RDS featured a central region where disparity varied (updated at the display’s frame rate, either 100 Hz or 60 Hz) and a surrounding annulus (1° wide, shown at 0° disparity). The size and position of these elements were determined by the collective receptive field of the recorded neural units, with an average RDS center size of 3.6° and a mean stimulus eccentricity of 6.3°. On any given stimulus frame, all dots in the center shared the same disparity, though this disparity could change from frame to frame. Signal disparities (always one near and one far value per session) were chosen to broadly align with the disparity selectivity of most recorded units.

Signal frames were randomly mixed with "noise" frames. The disparity for noise frames was drawn from a uniform distribution of typically nine discrete, equally spaced values (symmetrical around 0° disparity; common values included -0.4° to 0.4° in 0.1° increments), encompassing both the near and far signal disparities. Signal strength, expressed as a signed percentage (negative for near signal trials, positive for far signal trials), was determined by the proportion of signal frames to noise frames and served to adjust the task’s difficulty. For instance, -10% signal meant that for 10% of the stimulus frames (randomly distributed over time), the central area had the near signal disparity, while the remaining 90% of frames had disparities drawn from the noise distribution. In 0% signal ("no-signal") trials, all frames originated from the noise distribution, rendering the correct choice undefined; monkeys were randomly rewarded on 50% of these trials. The target icons were slightly smaller than the main stimuli and always presented with 100% near and far signal.

### Receptive field positions

We measured a horizontal and vertical response profile for each unit separately (Seillier et al., 2017). These reflect the one-dimensional receptive fields examined along the horizontal axis (using an elongated vertical stimulus, typically a low-spatial frequency sinusoidal luminance grating 0.3–0.5° by 3–5°) or vertical axis (using a horizontally elongated stimulus). These were fit by independent Gaussian functions. Those fits were required to explain at least 70% of the variance (see: Ǫuinn et al., 2021).

## Supplementary Results

### Microsaccades show a weak directional bias towards attended locations

If microsaccades play a causal role in attentional modulation, they would be expected to be directed toward the attended stimulus more frequently during the fixation period. To test this, we analyzed microsaccades recorded while monkeys performed the disparity discrimination task, focusing on the frequency of direction relative to the attended stimulus location.

We found that, even when spatial attention and motor planning were dissociated, microsaccades exhibited a weak but reliable bias toward the attended location. The proportions were nearly balanced (52.9% toward, 47.1% away), yet both monkeys made significantly more microsaccades towards the attended location (M1: mean = 52.0%, 95% CI = [51.5%, 52.6%], t(69) = 7.22, p<0.001; M2: mean = 53.0%, 95% CI = [51.8%, 54.1%], t(31) = 4.38, p<0.001). A similar distribution was observed when considering only the first microsaccade within each trial, indicating that the absence of a strong directional bias was not driven by later events during stimulus viewing (Fig. S1B-C).

### Microsaccade direction cannot explain attentional modulation of neural activity

Even in the absence of a strong directional bias, it remains possible that microsaccades influence neural activity differently depending on whether they are directed toward or away from the attended stimulus (e.g., Yu et al., 2022). To test this possibility, we examined neural activity time-locked to microsaccade onset, separately for microsaccades directed toward and away from the attended stimulus, and for trials in which the attended stimulus was either inside or outside the neuron’s receptive field. If microsaccades gated attentional modulation, neural responses would be expected to show enhanced activity following microsaccades toward the attended stimulus, with reduced or absent modulation following microsaccades in the opposite direction.

We found that attentional modulation was robust and highly similar across microsaccade directions. Neural responses exhibited comparable attention-related increases following microsaccades directed toward and away from the attended stimulus, both in magnitude and time course (Fig. S1D-E). No systematic interaction between microsaccade direction and attentional condition was observed across the population.

These results indicate that microsaccades do not gate or trigger attentional modulation of neuronal activity in V2 and V3/V3a. Rather, attention-related changes in firing rate persist independently of microsaccade direction, suggesting that motor-related eye movement signals and attentional state contribute separately to neural responses in mid visual cortex.

---

**Figure S1.**
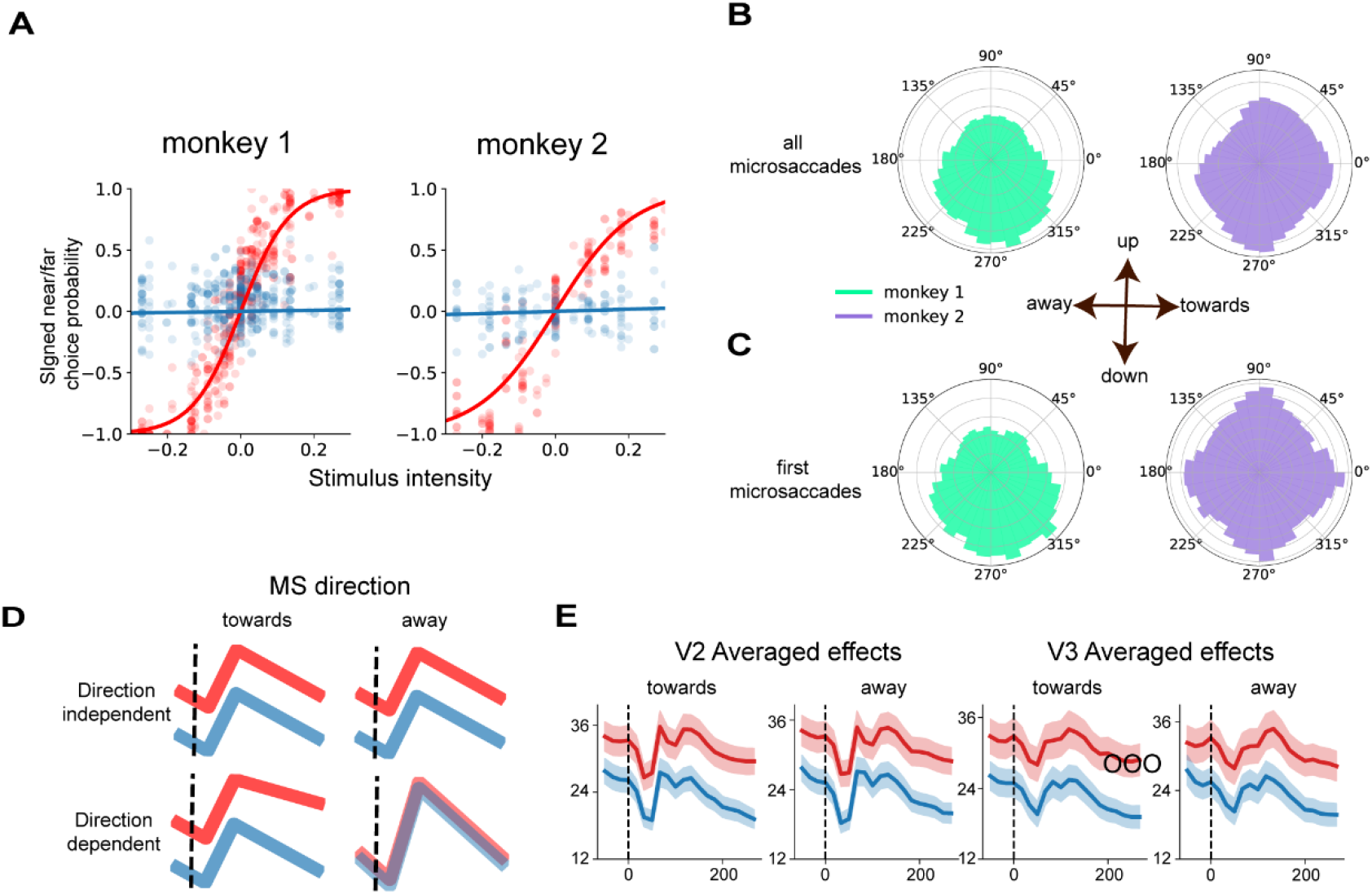
Behavioral performance and microsaccades. **A.** Psychometric functions showing *near* vs. *far* signed choice probability as a function of disparity signal strength (%), for *cued* (red) and *uncued* (blue) stimuli. Negative values indicate near disparities; positive values indicate far disparities. B-C. Because choice targets are decoupled from *cued* and *uncued* stimuli, the distributions of microsaccade directions are balanced across *toward* and *away* conditions for both monkeys. D. Hypothesis space for microsaccade-related neural activity. Top: attentional modulation is independent of microsaccade direction. Bottom: attentional modulation occurs only when microsaccades are directed *toward* the attended target. E. Microsaccade-triggered average activity as a function of attention and direction, indicating that microsaccade-related modulation is independent of microsaccade direction. Shaded areas represent ±2 SEM.

**Figure S2.**
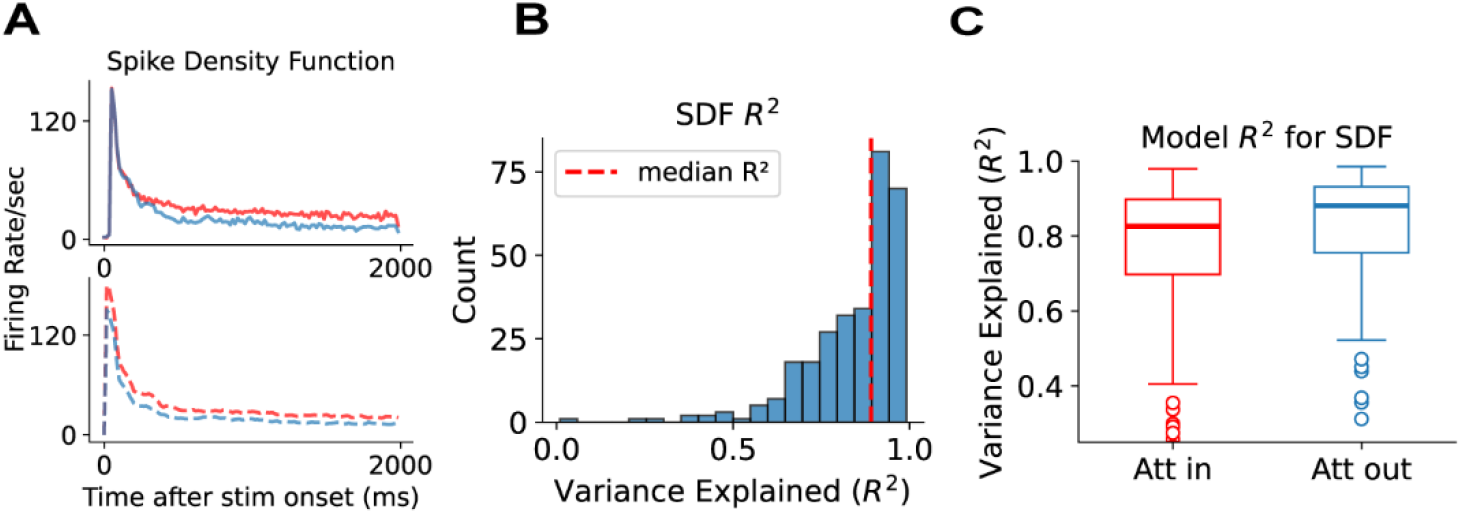
Building and validating the reference model. **A.** Spike-density function (SDF) variance explained by the reference model. Example unit: stimulus-triggered spike-density function (top) compared to model prediction (bottom). **B.** Histogram of SDF variance explained across all units by the reference model. **C.** Variance explained (R²) comparison across attentional conditions by the reference model. Boxes show interquartile range (IǪR); horizontal line indicates the median. Each point represents a single unit.

